# Disruptive changes in tissue microenvironment prime oncogenic processes at different stages of carcinogenesis in lung

**DOI:** 10.1101/2024.11.12.623197

**Authors:** Vignesh Venkat, Xiaoju Hu, Antara Biswas, Ankit Saxena, Jyoti Malhotra, Gregory Reidlinger, Subhajyoti De

## Abstract

Carcinogenesis is characterized not only by the uncontrolled growth of malignant cells but also by the disruption of the normal balance of cellular processes and intercellular interactions in the microenvironment that overcome the constraints of tissue homeostasis and support malignant growth. We profiled benign pulmonary dysplasia, carcinoma in situ, and invasive lung carcinomas at single-cell resolution to identify composite changes in cellular processes, signaling, and interactions among tumor-immune-stromal cells in the microenvironment in progressively advanced disease stages. We developed OncoTerrain, a hyperparameter-tuned model that captured synergistic multimodal signatures of an increasingly perturbed microenvironment in malignant disease stages and identified composite microenvironmental changes that supported cancer hallmarks. Key cancer-related changes in transcriptional states, cellular processes, and intercellular interactions involving immune, fibroblast, and stromal cell types preceded tumor initiation and were often synergistic. The microenvironment of increasingly malignant tissues was characterized by immune avoidance, ECM remodeling, and altered cell mobility. There were changes in cell states in fibroblasts, macrophages, and their inter-cellular interactions with other cell types, whereas T-cell activation occurred late. The in-situ carcinomas showed variations in the composite microenvironmental states that corroborated their pathology, which was not apparent at the genome level. A subset of those harbored populations of tumor and non-tumor cells with aggressive characteristics in some but not all aspects of hallmarks of carcinogenesis in the lung. We suspect that the variation in the coordination of microenvironmental cues may influence why some but not all in-situ tumors progress to the advanced stages.

## Introduction

The hallmarks of cancer^1^ not only involve uncontrolled clonal growth of malignant cells, but also dysregulation of ‘normal’ patterns of intercellular coordination among tumor, stromal, and immune cell types in the tissue microenvironment, disrupting the homeostatic constraints imposed by the design principles of multicellular life^2–4^. Abnormalities in macro- and microenvironmental characteristics often predate tumor initiation, as evident in the case of field cancerization and premalignant lesions – which has led to revisiting the seed and soil hypothesis of Paget^56^, originally developed for discussing metastatic tumors, and the theory of adaptive oncogenesis^7^. However, whether the molecular and phenotypic changes in tissue microenvironment follow coordinated patterns in progressively advanced disease, and whether those are directly related to the emergence of cancer-related complex hallmark processes remain to be systematically examined^1,3,8^. A major limitation in tracking cancer progression in humans is that many tumors arise asymptomatically and are detected late, such that it remains challenging to detect early changes in tumor progenitor cells as well as non-tumor cells in their microenvironment at the initial stages of carcinogenesis^1,8^. Age-associated hematologic malignancies and some types of solid tumors (e.g. skin malignancies) are notable exceptions, and some model systems provide critical insights in other cancers. Multiple investigator-initiated projects and large-scale consortia such as the Precancer Atlas projects^9^ have also provided important insights into multiple types of major solid tumors.

Historically, lung cancer has a high incidence rate, owing to smoking, environmental exposure, and genetic susceptibility^10,11^. The nationwide lung cancer screening program has only modest accuracy, leading to unnecessary follow-up examinations and invasive procedures. Although clinical guidelines are in place to increase detection accuracy, the classification criteria typically include only the size and texture of the nodules and their macro-characteristics^12^ and do not typically include molecular signatures from the tumor microenvironment^12,13^. Abnormal pulmonary nodules are commonly detected in low-dose CT in older adults, but a vast majority of suspicious pulmonary nodules do not progress to malignant disease, such that concurrently improving early detection and minimizing over-diagnosis remain clinical priorities. Key genetic, transcriptomic, and microenvironmental alterations in the major subtypes of lung tumors and premalignant lesions have been mapped which indicate multi-level heterogeneity^13–17^ but to what extent do changes in multiple different aspects of tumor cells and their microenvironment promote progression to advanced tumors was not examined. This motivated us to examine changes in tissue microenvironment at multiple levels across different stages of cancer development in the lung. Nationwide screening programs for high-risk individuals have banked a rich collection of malignant tissues as well as suspicious pulmonary nodules and other types of benign lesions^9,13^. Taking advantage of such resources, in this study, we profile biospecimens from benign pulmonary lesions, carcinoma in situ, locally advanced and metastatic carcinomas at single-cell resolution to assess multimodal and multiscale alterations in cellular processes and intercellular interactions involving tumor and non-tumor cells in tumor microenvironment in different stages of cancer progression (**Figure 1A**).

**Figure 1:**
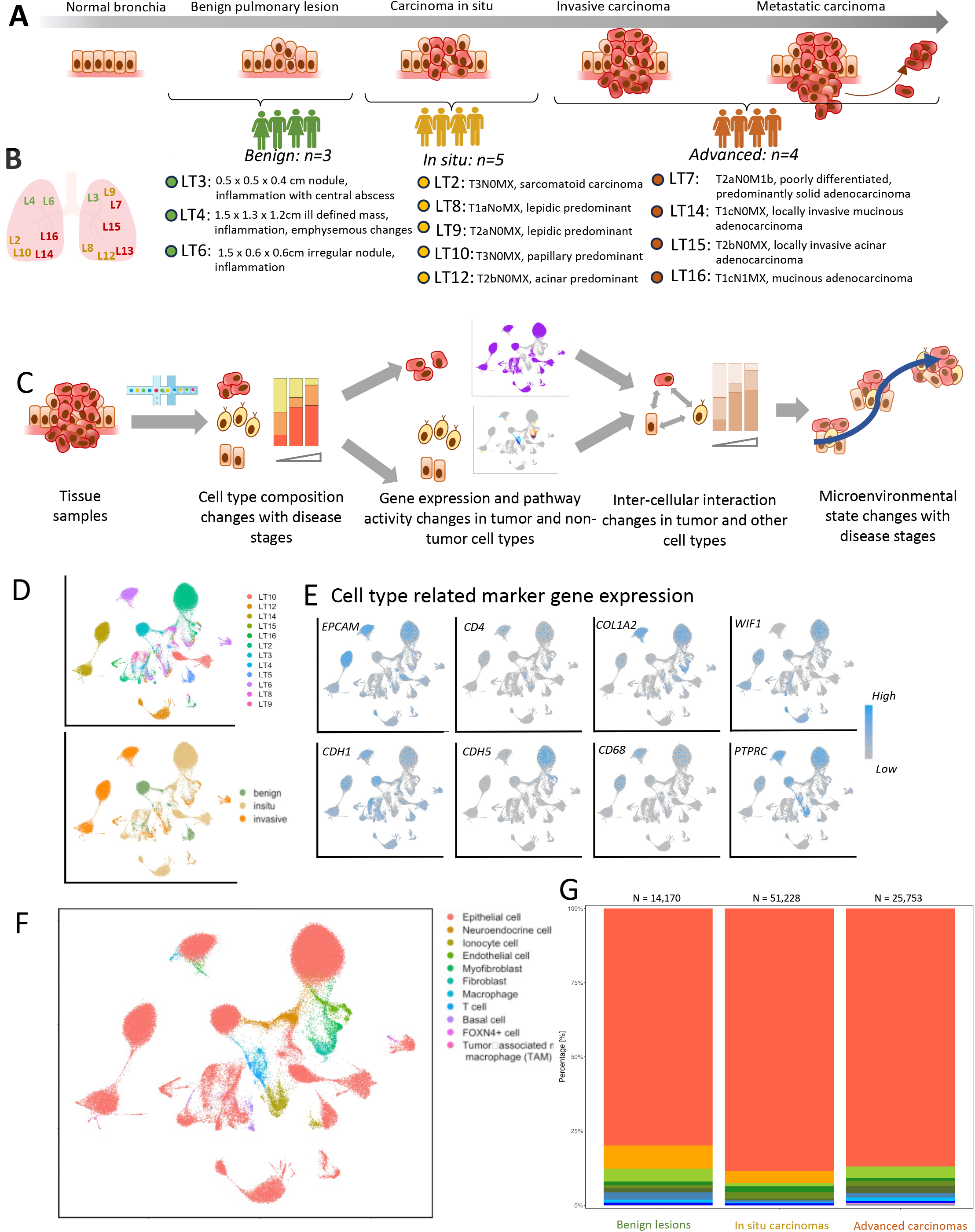
Overview of the study cohort and tissue composition. A) Profiling transcriptomic landscapes of the normal bronchia, benign lesions, in-situ carcinomas, invasive carcinomas, and metastatic carcinomas B) Location of sample and diagnosis from histopathologist C) Outline of study: Profiling the tissues, Cell type composition characterization with the onset of tumorigenesis, analyze cell-cell interactions, and utilize ML models to identify microenvironmental changes (D) Coloring the UMAP by sample ID and cancer stage (E) Coloring the UMAP by Cell Type Markers (F) Coloring the UMAP by Cell Type (G) Cell Type composition in the samples grouped by disease stages – i.e. benign lesions, in situ tumors, and invasion carcinomas.

## Results

### Cohort selection and transcriptomic profiling

The cohort for this study consisted of benign pulmonary lesions (n= 3; LT3, LT4, LT6), in-situ carcinoma (n = 5; LT2, LT8, LT9, LT10, LT12), locally invasive carcinoma (n = 4; LT7, LT14, LT15, LT16), and those with metastasis (**Figure 1B**), obtained from de-identified patients obtained under an IRB-approved protocol. The samples had representations from all lobes of the lung and different pathological subtypes. The benign pulmonary lesions were heterogeneous, and often accompanied by granulomatous inflammation. The pathological report for LT3 signified necrotizing granulomatous inflammation in the right upper lobe of the lung along with a hyalinized calcified granulomas lymph node. LT4 was noted to have scars, non-necrotizing granulomatous inflammation, and emphysematous changes. LT6 had a left pulmonary nodule, and a GMS stain showed histoplasma, and granulomatous inflammation in the level 5 lymph node. We chose to include proportionally more in situ carcinoma samples, which represent early stages of cancerous progression, and are often difficult to classify for clinical management. The in-situ carcinoma samples were diverse in their pathological annotation. LT2 had a left thoracotomy along with a lobectomy on the left lower lobe chest wall and was diagnosed with a chest wall tumor. LT10 was diagnosed as a papillary predominant adenocarcinoma in the lower, left lobe of the lung. LT15 was determined to have a right/middle lung invasive adenocarcinoma with an acinar subtype that exhibited a predominant cribriform pattern (4.1 cm diameter). LT12 showed signs of a papillary dominant, infiltrating adenocarcinoma (6 cm diameter). Some in situ carcinoma samples were of mixed histology. LT8 showed an infiltrating adenocarcinoma of the lung (2 cm in diameter), with lepidic (50%), papillary (5%), and acinar (45%) pattern characteristics. Some samples had mixed histology. LT7 showed the right upper lobe differentiated adenocarcinoma with a predominant solid pattern with focal acinar areas and tumor extensions into visceral pleura. LT14 was diagnosed with invasive mucin-producing adenocarcinoma, with solid, acinar, and micropapillary subtypes being predominant, and LT16 was diagnosed with a left lower lobe lung, moderately differentiated adenocarcinoma (3.0 cm diameter).

### Inference about cell type composition in the microenvironment

We performed single nuclei sequencing on the biospecimen using the 10X Chromium platform to characterize cellular transcriptome in the samples and used that to infer cell type composition, cellular processes, and intercellular interactions across different stages of disease progression (**Figure 1C**, see **Methods** for details). We processed the single nuclei transcriptomic data (see **Methods** for details) and identified stable cell type clusters based on their transcriptomic profiles (**Figure 1D**), which were then annotated using known cell type-specific marker gene sets (**Figure 1E-1F**). We showcase it by annotating the cellular barcodes with known markers for epithelial cells (e.g. EPCAM, CDH1), lung alveolar cells (SFTPC, WIF1), immune cell types including lymphoid and myeloid cell types (PTPRC, CD4, CD8), and fibroblasts (COL1A2, FBLN1; **Figure 1E**) that are a major component of normal and malignant lung tissues^13,18,19^, while additional markers are reported in cell type-specific contexts later. Overall, epithelial cells were the predominant cell types, as expected, and the major stromal (endothelial and basal cells) immune cells and fibroblasts typical of lung histology were also present in all samples (**Figure 1G**). Further annotation of the transcriptomic clusters by their sample of origin and the stages of cancer development indicated cell-type dependent differences (**Figure 1D-1F**). The epithelial cells, a majority of which are potentially neoplastic, especially in the malignant samples, formed multiple transcriptomic clusters that were predominantly segregated by the sample ID, and those of the advanced tumors were more different from one another; this pattern observed elsewhere as well might be due to accumulation of extensive genomic alterations unique to each tumor^20^. In contrast, the cell type clusters of non-tumor cell types typically had representations from multiple samples, indicating more nuanced inter-patient variation.

Epithelial cell fraction varied from 61%-90% in the benign samples, to 37.5%-92.4% in the in-situ samples, and 77.2%-91.6% in the invasive or distant samples, and the vast majority of those, especially in the advanced tumors were likely of malignant progenitor origin (**Figure 1G**). The proportions of immune cell types were relatively higher in the benign samples, where the T cell fraction for benign samples varied from 0.04%-1% to 0%-0.8% in the in-situ samples, and 0.3%-2% in the invasive samples. The macrophage cell fraction varied from 1%-4% in the benign samples, 0.2%-8.89% in the in-situ group, and 0.08%-3% in the invasive group. Relatively high immune cell presence might be due to inflammation noted for these samples. In contrast, in situ, tumors had relatively less immune cell infiltration, representing a relatively immune-cold environment, which reversed again in the advanced tumors that had higher proportions of different types of immune cells (**Figure 1G**). The proportions of fibroblasts increased from benign lesions and in situ tumors to advanced tumors, which is consistent with fibrosis in large or advanced carcinomas. Benign samples have proportions between 0.1%-7.7%, in-situ samples have proportions between 1-2.9%, and invasive samples have proportions between 0.8%-8.4%. The fibroblasts in advanced carcinomas had a prominent expression of known markers such as COL1A2 and FBLN1 and could be further segregated into those with myofibroblast characteristics. COL1A2 expression was only found in 48.9% of benign fibroblast and myofibroblasts, 48.8% of in-situ fibroblasts and myofibroblasts, however, was found in 80.5% of fibroblast cells in invasive tumors. FBLN1 expression was found in 15.6% of benign fibroblasts/myofibroblasts, 41.2% of in-situ fibroblasts/myofibroblasts, and 28% of invasive fibroblasts/myofibroblasts (**Figure 1G**). The extent of heterogeneity and richness in cell type composition, measured using Shannon’s and Simpson’s indices showed sample- and disease-stage-level differences, where benign Shannon indices ranged from 0.47 to 1.397 for the benign samples, 0.48 to 1.16 for the in-situ samples, and 0.16 to 1.27 for invasive samples. The Simpson’s indices ranged from 0.83 to 1.65 for the benign samples, 0.63 to 1.68 for the in-situ samples, and 1.09 to 1.91 for the invasive samples. A lower Shannon and Simpson’s index indicates lower cell type diversity within a sample, while a higher index signifies higher diversity with a more even cell type distribution. Heterogeneity in cell type proportion in the samples within and across the disease stages suggests disruptions of the normal balance of tissue composition with malignancy in the lung, like that observed elsewhere^16,17,19,21^. Variations in cell subtype-specific and cellular process-specific markers within cell-type clusters within and between patients motivated a more detailed, cell-type-specific, and context-dependent analysis^22^ (**Figure 1E-1F**).

### Functional heterogeneity in epithelial cell populations

Epithelial cells showed extensive intra- and inter-sample transcriptomic heterogeneity in the cohort. The epithelial cell transcriptomic clusters were typically segregated by sample IDs, and transcriptomes of the epithelial cell clusters in advanced tumors were more different from one another than those in the benign lesions - suggesting increasing between-sample transcriptomic heterogeneity with the disease stages (**Figure 2A**).

**Figure 2:**
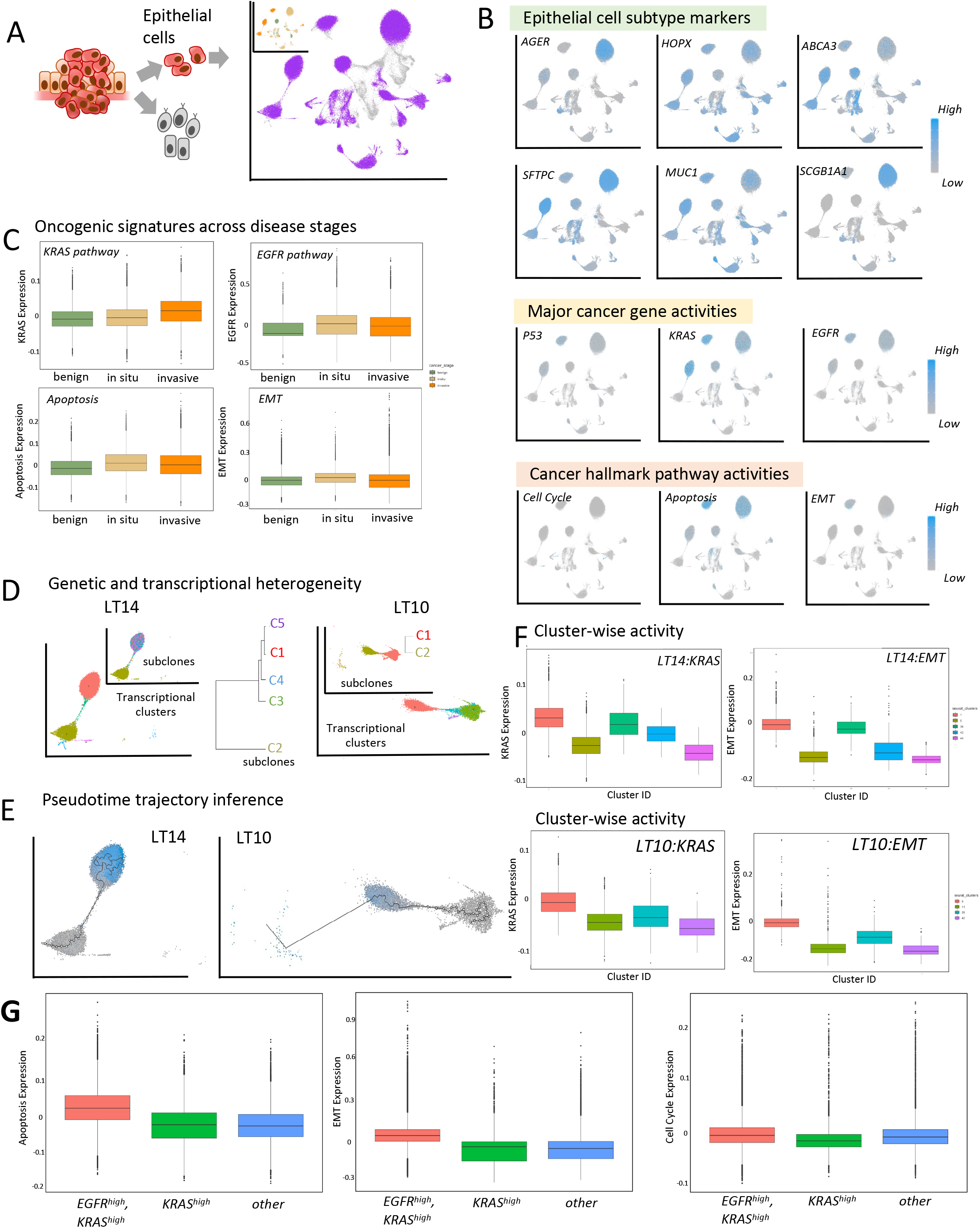
Functional heterogeneity of the epithelial cell population. A) The clusters of epithelial cell populations are examined for functional characteristics and cancer hallmarks. B) Annotating the epithelial cell clusters on the UMAP based on epithelial cell subtype markers, activities of pathways related to known cancer driver genes, and cancer hallmarks. (C) Boxplots showing changes in the activities of selected pathways related to cancer genes and hallmarks across different disease stages. Pathways associated with KRAS and EGFR activities, apoptosis, and epithelial-mesenchymal transition are shown as examples. D) Relationship between subclones with distinct copy number differences and transcriptional heterogeneity in advanced carcinomas in the lung. Sample LT14 and LT10 were shown as examples. E) Inferred transcriptional trajectories within the epithelial cell populations in LT14 and LT10 show a gradient in transcriptional changes in pseudotime. F) Significant differences in selected cancer hallmark pathway activities between the subclonal clusters in LT14 and LT10. All between-cluster differences are significant (Mann Whitney U test, FDR adjusted p-value < e-02). F) Significant differences in selected cancer hallmark pathway activities between *EGFR*^*high*^*KRAS*^*high*^, *KRAS*^*high*^, and *other* samples in the cohort. All between-cluster differences are significant (Mann Whitney U test, FDR adjusted p-value < e-02).

Annotation by lung cell subtype-specific markers, as well as those associated with oncogenic processes and cancer hallmarks^13,16,21–23^ revealed functional significance of the observed heterogeneity. Most of the epithelial cell subpopulations were primarily SLC16A7+, although some samples had populations of epithelial cells expressing additional cell subtype markers consistent with differences in histologies. SLC16A7 is an important biomarker as it indicates the tumor microenvironment is acidic, enabling cancer cell survival and immune evasion while also supporting the metabolic needs of rapidly proliferating tumor cells. For instance, LT7 which had a mixed histology with predominantly solid characteristics had cell clusters annotated with secretory, SLC16A7+, and ciliated cell type markers (**Figure 2B**). LT2 which is a sarcomatoid carcinoma had representations from SLC16A7+, ciliated, and club cells. LT14 and LT16, both of which are advanced carcinomas, had prominently elevated expression of the EGFR pathway (**Figure 2C)**. Interestingly, LT6, which is a benign granuloma also had high EGFR expression. We also noted that LT7, LT9, and LT12 had high *KRAS* expression, suggesting RAS-driven tumorigenesis. Notably, LT7 is an acinar carcinoma, while LT12 is of papillary subtype - both are known to have frequent KRAS mutations^24^. We also note that the RAS-driven samples have P53 downregulation, leading to upregulation of the cyclins and furthering cell proliferation^25^.

Advanced tumors also showed high intra-tumor transcriptomic heterogeneity in epithelial cell populations, which often formed multiple transcriptomic clusters with prominent differences in oncogenic driver and hallmark pathway activities^15^. Annotation of tumor subclones using inferCNV suggested that in the advanced tumors the transcriptomic clusters with different functional characteristics were usually associated with different subclones (**Figure 2D-2E**). We noted that in LT14, cluster 1 and cluster 36 were related to C1, C3, C4, C5, and cluster 5 was related to C2 (**Figure 2D-2E-2F**). We further investigated the KRAS and the Epithelial-Mesenchymal Transition (EMT) signature differences between the clusters to correlate genomic drivers with transcriptomic changes. We observed that KRAS pathway module score was higher in the transcriptomic cluster 1 (*μ =* 0.031) and cluster 36 (*μ =* 0.017), in stark contrast to cluster 5 (*μ =* −0.0302), cluster 43 (*μ =* −0.0066), cluster 44 (*μ =* −0.0465). We also observed the EMT hallmark pathway module score was higher in the transcriptomic cluster 1(*μ =* −0.0104) and cluster 36 (*μ =* −0.0269), which differs from cluster 5 (*μ =* −0.151), 43 (*μ =* −0.111), and 44 (*μ =* −0.162). Given the elevated activity of KRAS and EGFR pathways in the cluster C1 (**Figure 2B-2D-2E**), it is plausible that the clonal augmentation is RAS-driven distinguishing it from the counterpart clusters 5 or sub-clonal population C2^26^. The uptick in EMT signaling (**Figure 2C**) could further suggest a potential for cellular plasticity and cell state dynamics with metastatic potential in this cluster compared to the other clusters in this tumor^27^. We also conducted a similar analysis for LT10, where cluster 6 mapped to C1 and clusters 11, 28, 40 to C2, respectively (**Figure 2D-2E**). The clusters 6 (*μ =* −0.00930) and 28 (*μ =* −0.0376) had higher KRAS pathway activity in contrast to the clusters 11 (*μ =* −0.0474) and 40 (*μ =* −0.0585). Furthermore, the cluster 6 (*μ =* −0.0284) and 28 (*μ =* −0.0981) had greater activation of the EMT pathway in contrast to cluster 11 (*μ =* −0.147) and 40 (*μ =* −0.157). Many of these tumors were also annotated to have mixed histologies with characteristics that are in line with the observed intra-tumor functional transcriptomic differences. These results suggest that many advanced tumors show extensive intra-tumor functional transcriptional heterogeneity that correlates with intra-tumor sub-clonal differences (**Figure 2D-E**). However, there were other intra-tumor transcriptomic variations in the pathway signatures and pathological attributes beyond that captured by sub-clonal genetic differences alone, underscoring the significance of non-genetic functional variations in non-small cell lung cancer as reported previously^13,28−30^.

Signatures of several cancer hallmark pathways such as cell cycle and apoptosis were more prominent in advanced tumors compared to early-stage lesions. We grouped the samples as *EGFR*^*high*^*KRAS*^*high*^, *EGFR*^*high*^, *KRAS*^*high*^, and *Others* based on the activity levels of EGFR and KRAS pathways in the epithelial cell populations therein (**Figure 2G**; see **Methods** for details) The pathway-level activities may not necessarily involve genetic changes in these driver genes. We noted that the epithelial cells in the *EGFR*^*high*^*KRAS*^*high*^ samples had a significantly higher mean apoptosis hallmark pathway module score (*μ =* 0.0338) than the epithelial cells from the samples categorized as *KRAS*^*high*^ (*μ =* −0.0167) and *Others* (*μ =* −0.0165; FDR adjusted p-value <e-02). Elevated cell cycle and apoptosis pathway signatures are likely consequences of sustained growth signal, nutrient availability and replication stress in cycling cells in rapidly growing malignant tumors and pairwise correlated activities. Collectively these observations suggest elevated growth dynamics in the tumors driven by activation of one or more oncogenic driver pathways^16^.

Lastly, we performed a GSEA analysis^31^ to assess which cancer Hallmark pathways are differentially up- or down regulated in the epithelial cells in the advanced carcinomas compared to that in the benign lesions. We observed that MYC pathway, which regulates many genes involved in cell growth, proliferation, and metabolism, and often associated with aggressive malignant phenotypes, was significantly upregulated in advanced tumors, in line with that Dang et al suggests^32^ (*ρ =* 0.794, *p =* 0.0035). Genes in the Reactive Oxygen Species pathway, which are associated with cell stress, DNA damage, and inflammation, and likely related to elevated cell cycle activity was upregulated as well (*ρ =* 0.931 (pathway NES), *p =* 0.012). Successful tumor growth requires immune avoidance; we noted a significant upregulation of the Allograft rejection hallmark pathway in the advanced tumors (*ρ =* 0.828, *p =* 5.55 × 10^−6^) possibly reflecting the tumor’s ability to evade immune detection or the presence of an immune-related rejection mechanism^33^. This is notable considering increased abundance of immune cell types and heterogeneity in some of the immune cell type markers (e.g. IRF4) associated with ineffective immune surveillance in advanced tumors^34^.

### Functional heterogeneity in immune and stromal cell populations

The transcriptomic clusters of immune cells, fibroblasts, and other stromal cells in the cohort were segregated by cell types, and those cell type clusters typically had representations from multiple samples (**Figure 3A**) - which suggests modest inter-individual variations that contrasts the high inter-sample differences observed for the epithelial cell clusters. Systematic variation of some lung-cancer-associated marker genes and cellular processes within the cell type-specific transcriptomic clusters^16,18,21,35^ motivated to ask whether some of the differences are likely to be due to presence of cell subtypes and/or cell states therein (**Figure 3A-3B**).

**Figure 3:**
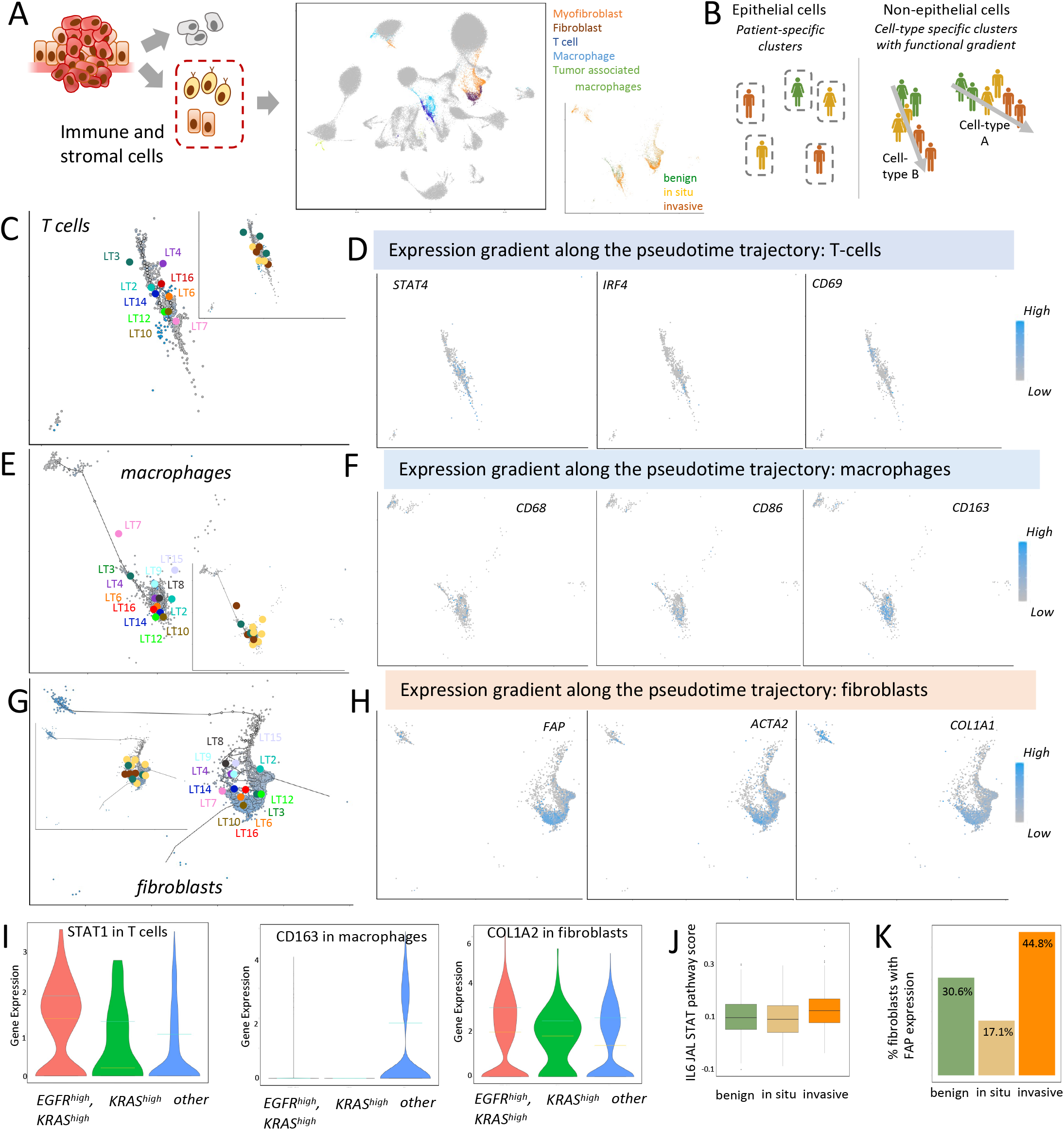
Functional heterogeneity of the non-tumor cell populations in the tissue microenvironment in the samples. A) UMAP projections of the barcodes grouped by cell types and disease stages. Non-epithelial cells are shown in different colors, while epithelial cells are in grey. Within respective non-epithelial cell types, the barcodes from different samples show considerable overlap and stratification by disease stages. B) A schematic representation showing functional annotation and heterogeneity of the populations of non-epithelial cell types in the samples across different disease stages. C) Inferred transcriptional trajectories within the T cells in pseudotime and D) transcriptional gradient of selected genes along that transcriptional trajectory. E) Inferred transcriptional trajectories within the macrophages in pseudotime and F) transcriptional gradient of selected genes along that transcriptional trajectory. G) Inferred transcriptional trajectories within the fibroblasts in pseudotime and H) transcriptional gradient of selected genes along that transcriptional trajectory. I) Box plots showing differences in cell-type-specific COL1A2, STAT1, and CD163 expression in the samples that are EGFR^high^KRAS^high^, KRAS^high^, and *Other*. COL1A2, STAT1, and CD163 expression were shown in fibroblasts, T cells, and macrophages, respectively. J) Significant differences in the IL6-STAT pathway scores in the T cell populations between benign, in situ, and invasive carcinomas. Mann Whitney U test, FDR adjusted p-values < e-02. K) Significant differences in % fibroblast cells with detectable FAP expression between benign, in situ, and invasive carcinomas.

The T cells from different samples formed a transcriptomic cluster, which showed structured transcriptomic heterogeneity therein with a principal axis of variation captured by a Monocle^36^ trajectory projection through the cluster (**Figure 3C)**. Annotation by sample ID and marker genes showed both intra- and inter-sample transcriptomic heterogeneity also along the principal axes of variation; interestingly, the trajectory was functionally relevant, correlated with expression gradient of multiple T cell marker genes (**Figure 3D**; IRF4, CD69, STAT4), and the transcriptomic gradient reflected progressively advanced stages of malignancy.

Furthermore, a subset of the T cells along the trajectory carried the characteristics of regulatory T cell type, while others were activated type, and the latter was more common in the advanced tumors. For instance, T cells of the benign and early-stage tumors such as LT3, LT4 and LT16 were clustered together and collectively presented high CD69 expression, which is a marker of Treg cell differentiation as well as that of precursor and mature resident memory T cells localized in peripheral tissues^37^. In contrast, that of the LT6, LT14, and LT2 overlapped and showed an increase in the STAT4 expression. The T cells of malignant lung carcinomas such as LT10, LT7, and LT12 (**Figure 3C**) were clustered together and displayed an increasing trend in STAT4 and IRF4 expression. STAT4 is upregulated by TCR stimulation in response to cytokines, and may regulate PD1 expression^38^, while IRF4 is a marker of T helper cell differentiation, and its upregulation is often observed in the context T cell exhaustion, such that their collective expression pattern may suggest an immunosuppressive environment^34^.

Similarly, macrophages also showed systematic variation in transcriptomic profiles between patients (**Figure 3E**), with relatively more discrete clusters with M1 and M2 subtype characteristics – which suggested dynamics between M1 and M2 type Macrophages in progressively advanced stages of cancer^39^. The benign lesions and early-stage tumors such as LT4, LT2, LT9, LT3, LT8 were tightly clustered and showcased a higher expression of CD86 and CD68 (**Figure 3F**). High expression of these markers is typically associated with M1 macrophages and suggest a pro-inflammatory state, where the macrophages are participating in immune surveillance or responding to inflammatory stimuli within the tissue^39^. In contrast, advanced tumors such as LT15 and LT7 showed more variability with their centroid’s position on the UMAP, which indicates a transition from M1 to M2 type macrophages. The macrophage cell populations of malignant tumors LT16, LT6, LT14, and LT10 were all tightly clustered, and showed high CD163 expression suggesting a prominent presence of M2 macrophages, which are typically involved in anti-inflammatory responses and tissue repair^39^. There might be similar variations in other immune cell types^21,30^, but the number of such cells was not sufficient in our cohort to warrant a systematic analysis.

Fibroblasts are key components of tissue microenvironment and show stage specific transcriptional plasticity^35^. Fibroblasts in our cohort formed functionally relevant subclusters (**Figure 3G**), a subset of which identify myofibroblast-like cells. FBLN1 showed an increase in expression in the myofibroblast and fibroblast cell populations, indicating an increase in glycoprotein secretion, responsible for components to the fibrillar extracellular matrix^40^. Monocle analysis indicated a similar axis of intra- and inter-patient variations across fibroblast subtypes that was also associated with the disease stage towards malignancy. The samples LT15, LT8, LT4, and LT9 were all closely clustered, and had an upregulation in COL1A1 (**Figure 3H**), which indicates an active role in tissue remodeling, wound healing, or fibrosis processes. In contrast, the fibroblast subcluster including LT6, LT10, LT16, LT3, and LT12 had an increase in ACTA2 and FAP expression (**Figure 3H**). FAP indicates a role in degrading extracellular matrix components and modulating the local tissue environment, which is critical for processes like fibrosis and tumor progression^41^. Other samples such as LT7, LT14, and LT2 had more diverse patterns; LT7 and LT14 showcased a higher expression of COL1A1, COL1A2, and ACTA2 (**Figure 1E; Figure 3H-3I)**. ACTA2 showed clear activation signals in the fibroblasts and showed a gradient from the myofibrotic cells to the fibroblasts in pseudotime, suggesting an increase in alpha-2 actin proteins, which are heavily involved in cell motility, structure, integrity, and intercellular signaling^42^. LT2 showcased a transcriptional gradient that lacked correlation with any major known cancer-related fibrosis markers except COL1A2 upregulation. The transcriptional trajectory patterns for fibroblasts and T cells suggest phenotypic plasticity and cell state continuum, and associated gene expression changes signify their relevance for remodeling of extracellular matrix and tissue microenvironment in general.

Grouping by the disease stage and EGFR/KRAS pathway status revealed driver-pathway related changes in the non-tumor cells in the microenvironment (**Figure 3I**). For instance, immune marker gene STAT1 was overexpressed in the T cell populations in the *EGFR*^*high*^*KRAS*^*high*^ (*Q*_1_ = 0.00000634, *Q*_2_ = 1.37, *Q*_3_ = 1.92) and *KRAS*^*high*^ (*Q*_1_ = 0.00000407, *Q*_2_ = 0.192, *Q*_3_ = 1.31) groups in contrast to the *Other* (*Q*_1_ = - 0.00000180, *Q*_2_ = 0.00000963, *Q*_3_ = 0.996) group. We also grouped the macrophages by the EGFR/KRAS pathway status of the tumors, and found the *Other* group had an upregulation in CD163 (*Q*_1_ = -0.00000479, *Q*_2_ = 0.00000388, *Q*_3_ = 1.85) in contrast to *KRAS*^*high*^ (*Q*_1_ = -0.00000177, *Q*_2_ = 0.00000410, *Q*_3_ = 0.00000512) or *EGFR*^*high*^*KRAS*^*high*^ (*Q*_1_ = -0.00000541, *Q*_2_ = 0.00000221, *Q*_3_ = 0.0000111). Similarly, fibroblast signatures showed an upregulation of COL1A2 in the *EGFR*^*high*^*KRAS*^*high*^ (*Q*_1_ = 0.00000462, *Q*_2_ = 1.91, *Q*_3_ = 2.96) and *KRAS*^*high*^ (*Q*_1_ = 0.0000180, *Q*_2_ = 1.74, *Q*_3_ = 2.39) groups in contrast to the Other (*Q*_1_ = 0.000000837, *Q*_2_ = 1.34, *Q*_3_ = 2.53) group. Other studies have shown that KRAS-mutated group is associated with lower immune infiltration^43^, lower expression of immune checkpoints, especially, a lower abundance of B cell, CD8+ T cell, dendritic cell, natural killer cell, and macrophage, higher abundance of neutrophil and endothelial cell, and that immunotherapy is less effective in patients with EGFR mutant non-small-cell lung cancer^44^. Based on the literature and our results, the trajectory-oriented gene signatures, abundance of immune cell (**Figure 3J)**, fibroblast subtypes in the samples (**Figure 3K)**, and the disease stage, suggest that both abundance and cell states of the non-tumor cell types change with tumorigenesis and altered oncogenic driver pathway activity.

### Functional heterogeneity in intercellular signaling

Heterogeneity in the relative abundance of cell types and their cellular processes, led us to examine the patterns of intercellular interactions involving epithelial, immune and stromal cells in the tumor microenvironment and changes thereof in progressively advanced stages of carcinogenesis in lung (**Figure 4A**). Firstly, we examined inter-cell type communication patterns of ligand-receptor pairs from the benign lesions, in situ tumors, and invasive carcinomas (**Figure 4B**). As an example, the benign samples had inferred interactions of multiple ligands - LAMB1, LAMA2, FN1, COL6A3, COL4A2 with the CD44 receptors, but such interactions weaned in the in-situ samples before returning slightly in the invasive samples. These proteins are important components of the basement membrane and interstitial matrix ^45^, and their ligand-receptor interactions likely suggest intact and structured tissue architecture and extracellular matrix maintenance in benign samples, which is disrupted in the malignant samples. In the in-situ samples, we observed significant ligand-receptor interactions between SLIT2 and ROBO1/ROBO2 receptors; although SLIT2-ROBO signaling typically helps to suppress tumor spread, disruptions in this pathway can contribute to tumor growth and metastasis^46^. On the other hand, we observed that multiple ligands - LAMA2, LAMB1, FN1, COL6A3, COL4A2/COL4A1, and COL1A2/COL1A1 had heterogeneous patterns of predicted ligand-receptor interactions with the CD44 receptors in the invasive samples. LAMB1/LAMA2, COL6A3, COL1A2 are known to interact with ITGA1+ITGB1, while FN1 ligands interact with ITGAV+ITGB1 receptors in the context of matrix maintenance and remodeling^47^. FN1 is a glycoprotein that plays a key role in cell adhesion and migration, and the genes involved in the FN1-CD44 pathway went through considerable increase in transcriptional activities from benign to the in-situ tumors^48^ (**Figure 4C-4D**). Furthermore, interaction of these ECM proteins with CD44 in an abnormal and heterogeneous manner could signify altered tumor cell motility and invasive potentials. Therefore, taken together, elevated activities of these ligand-receptor interactions may be suggestive of active ECM dynamics, which in turn may facilitate tumor cell invasion, migration, and metastasis. In addition, we also found increased expression of ligand-receptor gene pairs in the APP and CD74 pathway with tumor stages^49^ - which suggests a potential interplay between cellular signaling pathways and immune-related mechanisms that may be crucial to modulate tumor-immune system interactions during different stages of cancer progression.

**Figure 4:**
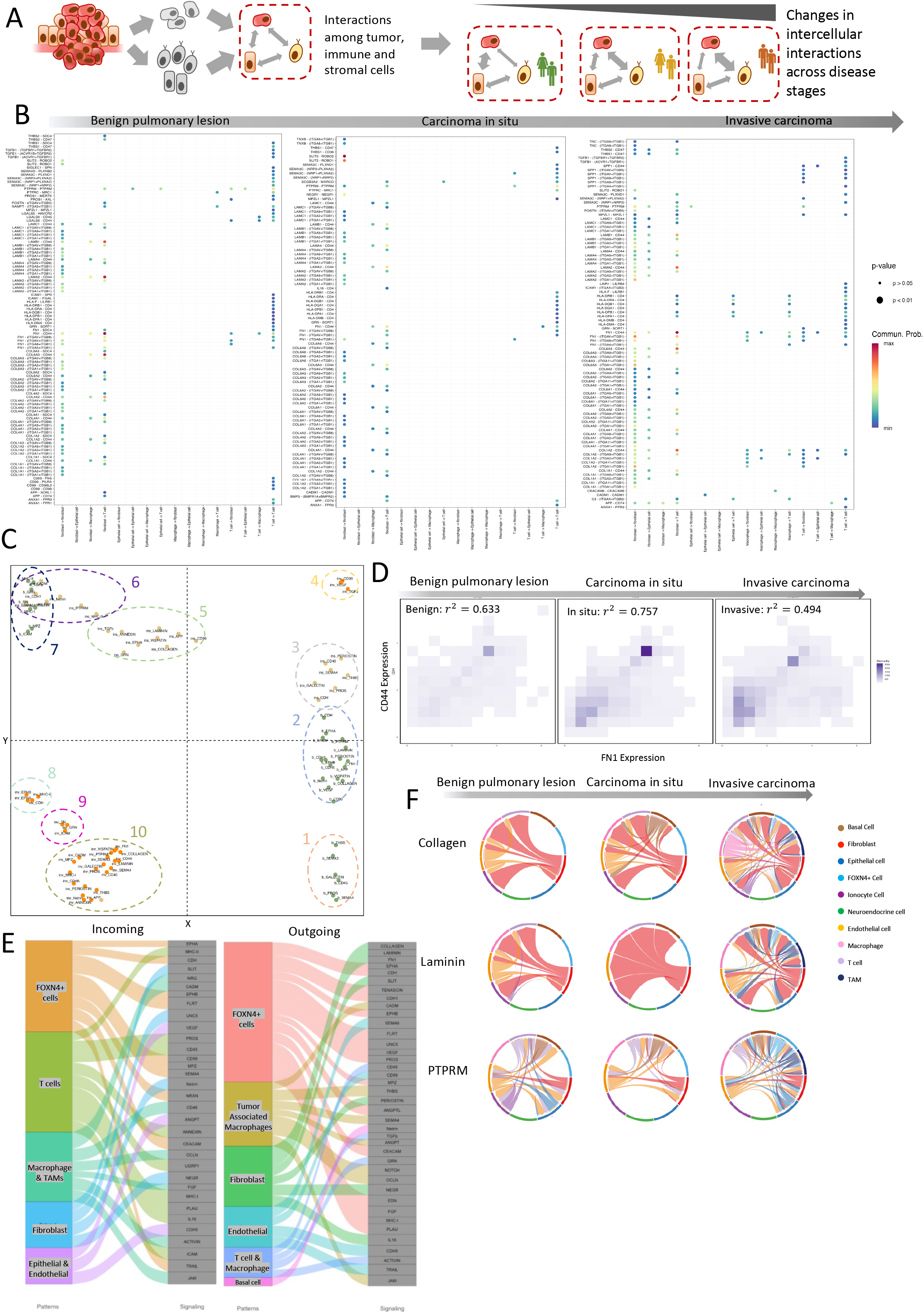
Functional heterogeneity in intercellular signaling between stroma, immune, and epithelium in the samples. A) Construct a CellChat object to discover ligand-receptor interactions between tumor, immune, and stroma and observe changes across disease stages. B) A dot plot showcasing the intensity of ligand-receptor interaction alterations through tumorigenesis, with the x-axis representing cell type-to-cell type interaction and the y-axis representing individual ligand-receptor pairs. C) A scatterplot visualizing signaling pathway interactions across disease stages, with each cluster representing closely interacting and/or related signaling pathways. (D) A 2D Heatmap showing correlation and expression of FN1-CD44 in benign, in-situ, and invasive samples. (E) A Sankey diagram showcased incoming and outgoing signaling pathways for each pattern, where the pattern was derived from the NMF of the cell chat object. We have mapped each of those patterns to a corresponding cell type. (F) Circos plots showcase collagen, laminin, and PTPRM signaling pathway augmentation through tumorigenesis, where we noted an increase in chaos consistent in many groups of intercellular interactions.

Next, we examined the ligand-receptor interactions to identify those interactions and associated signaling pathways that are sustained, down-regulated, or upregulated in the progressively advanced stages of carcinogenesis. A set of ligand-receptor co-expression activity patterns in the cohort involving: COLLAGEN, LAMININ, PTPRM, FN1, APP, SEMA3, MHC-II, THBS, Netrin, CADM, VEGF, ANNEXIN, CDH5, GRN, MPZ pathways-which are broadly related to cell adhesion was sustained across all disease stages. However, another set of ligand-receptor co-expression signatures involving inter-cell-type signaling (e.g. CD45, Galectin, GRN, SN) associated with lung development and tissue homeostasis, weaned progressively in the in-situ and advanced carcinomas. Many of these signaling pathways are also associated with immune surveillance and inflammatory response; e.g. Galectin and GRN signaling are often associated with inflammation and fibrosis ^50^; their activities in the benign samples may be partly related to the pulmonary inflammatory conditions reported, but we also found those in the immune-cold in situ tumors, indicating that their activities are not solely artefacts of inflammatory conditions reported in some benign lesions. In contrast, abnormal ligand-receptor interactions typically associated with signaling processes linked to malignancy, deregulation of normal development, and immune signaling (**Figure 4C**; TENASCIN, NOTCH, CD46, NRG, UNC5, COMPLEMENT, FLRT, SEMA6, CEACAM, NRXN, ANGPTL, ANGPT, OCLN, FGF, JAM, ncWNT, ACTIVIN, EDN, LAIR1, TRAIL, NEGR) became more prevalent across progressively advanced disease stages. Similar patterns of disruption of normal signaling and gain of malignancy-related signals have been observed elsewhere^16,51,52^, and reflect a key aspect of cancer as a disease of disrupted homeostatic constraints imposed by multicellularity^4^.

However, multitudes of ligand-receptor interactions showed complex overall patterns of underlying the cell-cell signaling and preferences for specific cell types, which motivated us to use non-negative matrix factorization (see **Methods** for details) to summarize the outgoing and incoming patterns of intercellular ligand-receptor co-expression and extract composite signatures of cell-cell signaling that show distinct characteristics across the cohort (**Figure 4E**). We identified 5 distinct incoming and 6 outgoing signatures (IS and OS), and all signaling pathways listed below had significance (*p* < 0.001), which show coherent cell-type preferences and association with cancer-related biological processes. Incoming Signature IS1 was dominated by FOXN4+ cells and involved EPHA, CDH, EPHB, FLRT, CD99, MPZ, SEMA4, NRXN, CEACAM, TRAIL signaling process, that have roles in cell adhesion. Signature IS2 involved T-cells and included genes such as MHC-II, PROS, CD45, ANNEXIN, UGRP1, MCHI, PLAU, IL16, ICAM), indicating immune surveillance. The signature IS3 involved macrophages and tumor associated macrophages and had a role in growth and differentiation. The signature IS4 was directed to fibroblasts and involved possibly neuroendocrine phenotype-related genes, while signature IS5 involving epithelial and endothelial cells were related to growth. For the outgoing signatures the outgoing signature OS1 was dominated by FOXN4+ cells, and included interactions involving EPHA, CDH, TENASCIN, CHD1, EPHB, FLRT, UNC5, VEGF, PROS, ANGPTL, CEACAM, EDN, GFG, MHC-I, PLAU, signature OS2 was comprised of interactions engaging CADM, THBS, SEMA4, ANGPT, NOTCH, OCLN, JAM and primarily involved tumor-associated macrophages, signature OS3 was made up of Collagen, Laminin, FN1, SLIT, Periostin, NEGR, IL16 interactions in fibroblasts, signature OS4 was made up of endothelial cells (and indicated SEMA6, CD99, CDH5, TRAIL), the outgoing signature OS5 primarily involved T-cells and macrophages (and indicated CD99, TGF-B, GRN, Activin), while OS6 involved Netrin in the basal cells.

When we compared the signatures across the disease stages, it was found that no signaling signature appeared or disappeared in its entirety across the disease stages. Rather, some interactions of the signatures weaned or grew in advanced malignancies, indicating a signaling rewiring rather than a modular process (**Figure 4C**). We found that the signaling involving Collagen, Laminin, and PTPRM show most remarkable changes in cell-cell interactions across the disease stages (**Figure 4F**). In benign samples, Collagen signaling was primarily involving intra-cell-type interactions involving fibroblasts as well as inter-cell-type interactions with, Macrophages (*Pscore =* 0.19), and T cells (*Pscore =* 0.87). But the in-situ samples showcased a marked degradation in the ‘normal’ patterns of cell-cell communication involving ligands from the fibroblasts to T cell (*Pscore =* 0.25) receptors. In the invasive samples, we see a wider dysregulation in the ligand-receptor interactions involving multiple cell types: ligands from fibroblasts were predicted to signal to the receptors from tumor-associated macrophages (*Pscore =* 2.10), epithelial cells (*Pscore =* 0.29), and T cells (*Pscore =* 0.75), indicating emergence of abnormal patterns of cell-cell interactions besides presumed loss of the normal patterns. Similarly, PTPRM and laminin signaling showed systematic changes in both the cell-cell pairing and intensity of cell-cell interactions across the disease stages. Apart from the epithelial cells, which were primarily of tumor-origin, especially in the malignant lesions, fibroblasts, macrophages, and T cells appeared to be the key players involved in the rewiring in malignant tumors.

Taken together, our data suggests that there are changes in cell-cell interactions in the tumor microenvironment in malignant and advanced disease stages, and key changes are related to the extracellular matrix remodeling through the CD44 receptor activation and the emergence of immune avoidance through Galectin and GRN signaling pathway deactivation. Furthermore, tumor cells likely play a key role in shaping its microenvironment through signals to the laminin and collagen structures in the lung. Taken together with findings from the prior section, we believe that the tumor is shaping the tumor immune microenvironment by switching macrophages from M1 to M2-like, avoiding direct T cell attack, and reshaping the ECM so that the normal tissue structure “melts” away.

### Attention-based model to map changes in cancer hallmarks through disease stages

Most of the cancer hallmarks are not tumor cell-specific, rather emerge due to tissue-level biological processes arising from complex cell-cell interactions in the tumor microenvironment, which can be conceived as a dynamic complex system. We argue that the transcriptional activities of the cells and cell-cell interactions therein manifest the microenvironmental state, and between-sample variations thereof reflect plasticity and transition of the microenvironmental state, especially when interpreted in light of the disease stages (**Figure 5A**). This motivated us to design a model that can incorporate cell-cell interactions, cell trajectories, and the microenvironmental heterogeneity of tumors to assess the potential for the tumor and non-tumor cells in a tumor to exhibit distinct characteristics. We developed OncoTerrain, the attention-based tree model (**Figure 5B**, see **Methods**) to re-examine the microenvironmental states of the individual in situ samples based on the cancer hallmark-related activities at a single cell resolution; while a majority of the tumor and non-tumor cells should conform to the pathologically assigned disease stage, we asked whether sub-populations of tumor and non-tumor cells in some samples could carry more aggressive hallmark characteristics compared to the overall pathological classification of those sample.

**Figure 5:**
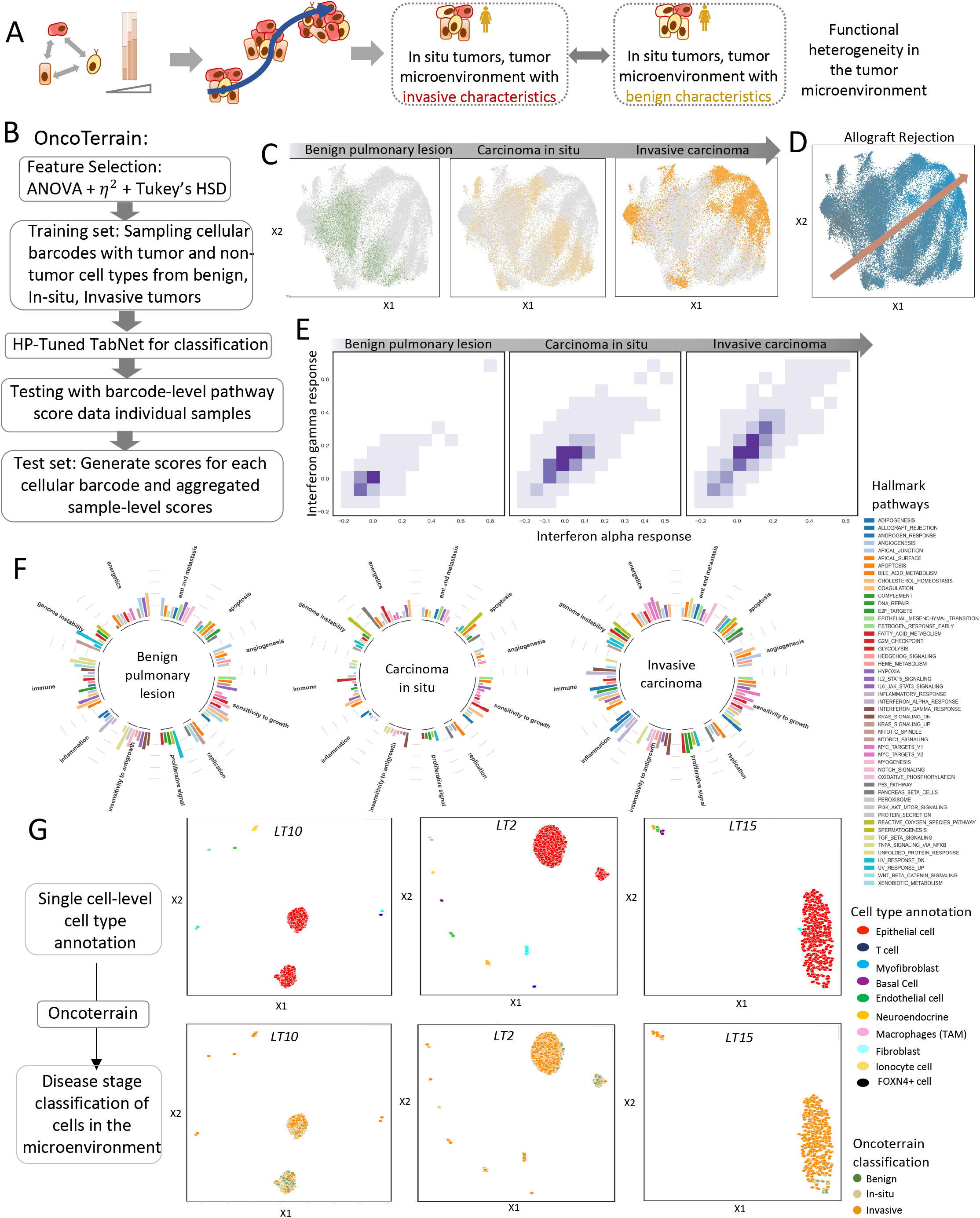
Utilizing OncoTerrain to map changes in cancer hallmarks through disease stages and relate pathway-pathway activity in the samples. A) A conceptual framework to examine the aggregated, cancer hallmark-related pathway-level changes in the tumor microenvironment across different disease stages. B) A reprojection of the barcodes using hallmark pathway module scores onto the UMAP hyperspace and we observe a far more homogeneous trajectory, albeit still heterogeneous, of tumor evolution. C) Coloring the UMAP with Allograft rejection pathway expression we notice a correlation between intensity of expression and stage of cancer. D) Outline workflow of OncoTerrain involving feature selection, curating a training set, tuning the hyperparameters, and inferencing upon individual samples. E) A 2D heatmap showing correlation and expression of Interferon Alpha-Interferon Gamma signaling during different disease stages. showed pathway characteristics of different disease stages. F) Circular bar graph showing hallmark pathway activity about the cancer hallmarks through stages of disease. G) Applying OncoTerrain on selected early-stage tumor samples -LT10 and LT2, and an advanced carcinoma SD15 where subpopulations of tumor and non-tumor cells showed pathway characteristics of different disease stages.

First, we calculated the cancer hallmark pathway scores (see **Methods**) and used the variance of the scores to reproject the cellular barcodes in the UMAP hyperspace. While individual barcodes retained their cell-type characteristics, cellular barcodes were more integrated when compared to the patterns observed at the cell-by-gene-level analysis (**Figure 1C-E**). We focused on their collective characteristics aggregated at the sample-level, which likely represents the activities of complex bioprocesses emerging from the tissue microenvironment state. When annotated by the disease stages of the sample-level cellular barcodes, we observed a gradual transition in the overall microenvironmental state from benign lesions to invasive carcinomas with certain in situ carcinomas clustering closer to benign lesions whereas others are trending towards the invasive cell population (**Figure 5C**). This contrasts the sample-wise discrete clustering patterns we observed for the epithelial cells and underscores emerging properties of the tumor microenvironment state considering cancer hallmarks, and its dynamics with the disease stages.

Upon examining stage-wise differences in the hallmark pathway signatures, we observed that Allograft Rejection (*p* < 0.001 ; *η =* 0.189) was the top pathway that showed significant augmentation through the axis of transition of microenvironmental state along the stages of tumorigenesis (**Figure 5D**). Allograft Rejection pathway in the context of carcinogenesis can be interpreted as heightened immune surveillance and/or evasion of anti-tumor immunity within the tumor microenvironment^53^. In addition, MYC targets (*p* < 0.001 ; *η =* 0.115), Interferon Alpha (*p* < 0.001; *η =* 0.111), and Interferon Gamma (*p* < 0.001; *η =* 0.108) were among the top pathways that also showed systematic variation along the axis of transition of microenvironmental states. The MYC oncogene regulates numerous genes involved in cell proliferation and its pathway activation in tumorigenesis underscores its role in promoting unchecked cell division and metabolic reprogramming^32,54^. Interferon Alpha^55^ is known for its immunomodulatory effects and the increased activity during tumorigenesis may reflect an attempt by the immune system to counteract tumor progression through immune-mediated cytotoxicity. This cytokine is critical for anti-tumor immunity, and its augmented signaling in tumorigenesis may suggest an active, though potentially ineffective, immune response aimed at controlling or eliminating tumor cells. We then examined whether tumorigenesis associated changes in these pathways in the tumor microenvironment are coordinated or independent. Using a 2D histogram we found that the activity of interferon alpha^55^ and gamma responses^56^ showed tight linear association (**Figure 5E**), which was also aligned with the axis of microenvironmental state transition during tumorigenesis; both pathways were upregulated in unison, suggesting a coordinated immune response involving both innate and adaptive immune defenses to combat malignant growth, and probable immune evasion by tumor cells.

We revisited the 10 original cancer hallmarks originally proposed by Hanahan and Weinberg^1^ in light of the observed changes in the microenvironmental state (**Figure 5F**). We first mapped the MSigDB cancer hallmark-related pathways to those hallmarks, allowing multi-mapping i.e. a pathway could be part of multiple hallmarks, when necessary, and examined to what extent the aggregated signatures attributed to the cancer hallmarks change across different stages of carcinogenesis in lung. Based on the collective activities of the pathways, we noted that some cancer hallmarks (e.g. genome instability, energetics, apoptosis) experience gradual increase through the tumor stages, while some others (e.g. insensitivity to antigrowth, sensitivity to growth) show more complex patterns, and yet some others show markedly augmented activities in the advanced carcinomas (e.g. immune, inflammation). Pathways therein showed mutually consistent patterns. For instance, within the cell cycle hallmark, the Reactive Oxygen Species pathway was upregulated^57^ and DNA damage response (UV response pathway; *p* < 0.001 ; *η =* 0.0643) was downregulated − which suggest increased DNA damage, replication errors, and impaired DNA damage response promoting rearranged genomes and genomic instability. Alongside ROS, we noted that P53-related cyclins were upregulated (i.e. P53 downregulation) was seen increasingly from benign to invasive samples (P53 pathway; *p* < 0.001 ; *η =* 0.0322). Similarly, upregulation of the energetics and apoptosis-related processes with the stages of tumorigenesis reflect the growing tumor’s need for energy to sustain rapid cell proliferation, alongside increased rates of programmed cell death as damaged or dysfunctional cells are cleared. In contrast, we noticed down regulation in the immune recognition and inflammation-related processes, which reflects the early tumor environment is primed for evading immune detection and beginning to suppress normal growth-inhibitory mechanisms^58^, creating a more favorable environment for tumor progression and becomes a dominating aspect of the microenvironment in advanced tumors. The relative changes in the hallmark-related activities reflected the variation in the coordination of microenvironmental cues in increasingly malignant disease stages.

The hallmark-level heterogeneity is especially relevant in the context of pathobiology of in situ tumors, since not all in situ tumors progress to become advanced disease. Thus, we applied OncoTerrain model, trained using a subset of the data, predominantly from the benign and advanced tumors, to test the single cell-level data in other ones, e.g. in-situ tumors (**Figure 5G**). Our model performed well (ROC-AUC = 0.95), and we tested for type I-II errors. We showcase LT10 - an in-situ, localized tumor of papillary predominant adenocarcinoma of size greater than 7.5cm, as a case study. Based on the hallmark pathway signatures, one of the clusters within the tumor cell populations therein emerged as predominantly early-stage, while the other cluster had primarily invasive characteristics. Clonal analysis indicated that the early-stage tumor cells are likely progenitor cells from the original clone C1 (**Figure 3E**). However, interestingly the stroma and immune cells belonged to more cohesive clusters with invasive characteristics - myofibroblast, fibroblast, macrophages, and T cells in this sample clustered with their counterparts in the invasive samples. Indeed, detailed histopathology report for this sample indicated mixed histology with papillary and lepidic pattern with likely intra-lobar metastasis (no lymph node or distal metastasis), supporting pathological manifestation of a likely invasive characteristics. We applied OncoTerrain to LT2 another early-stage tumors with complex histology. The model classified a subset of the cell populations with early-stage characteristics, while another cluster was trending towards an invasive characteristic. In this case study however, we noticed that the microenvironment was far less cohesive, with the fibroblasts and endothelial cells showing mixed transcriptomic profiles. The model inference suggested that the macrophages and T-cells were functionally heterogeneous – ranging from that like in the early-stage cancers to advanced-staged cancers. Therefore, we may consider LT10 and LT2 tumor to be associated with higher risk than their overall pathological classifications indicate. When testing OncoTerrain on an invasive tumor, LT15, our model identified cell populations therein with more late-stage characteristics in the microenvironment. Detailed pathological annotation indicated a locally invasive adenocarcinoma with mixed histology with acinar predominant characteristics, providing a rationale for the model observation (**Figure 5G**).

## Discussion

Cancer is as much a disease caused by genetic changes in the tumor cells, as it is a disease of a dysregulated tissue microenvironment^59^. Clonal growth of tumor cells is enabled by bypassing the constraints of multicellularity. In this work we analyzed the changes in the complex landscape of cellular processes and intercellular interactions involving epithelial, immune, and stromal cells from benign pulmonary lesions, carcinoma in situ, locally advanced and metastatic carcinomas at single cell resolution – which captured multiscale alteration in the tissue microenvironment states in different stages of cancer progression.

Genetic and transcriptomic heterogeneity in epithelial cells within and between tumors showcased functional heterogeneity. We observed that intra- and inter-tumor differences in oncogenic driver pathways had relevance for proliferation, EMT and other hallmarks, which can ultimately influence tumor evolution. While large scale changes in cellular transcriptome were often associated with sub-clonal heterogeneity (e.g. LT14 and LT10), our findings resonated with recent reports^13,29^ that non-genetic heterogeneity involving epithelial cell states was common and had functional relevance, which in turn could influence evolutionary dynamics of the tumors. Some key changes in cellular transcriptomes, cellular processes and intercellular interactions involving immune^34,43,44,48,58,60^, fibroblast^40–42^, and stromal cell-types were associated with the cancer hallmarks^1^ ; and some of those were synergistic, often preceded cancer initiation, while some others were established in the advanced stages of carcinogenesis. T cells, macrophages, and fibroblasts showed cell state dynamics within tumors and between disease stages, that are important for the plasticity and evolvability of the microenvironmental state. Tissue damage in the microenvironment in non-malignant tissues promotes inflammation and wound healing^61^, which may support cell proliferation and adaptive oncogenesis^7^. A subset of the benign lesions in our cohort had granulomatous inflammation in the vicinity, but no infection was identified. It is not known whether those are bystander events, but recent report suggests that inflammation in the tumor-adjacent lung can influence tumor-immune interactions and predict outcome^14^. Signatures of inflammation and immune-related processes were noted in all disease stages, albeit in situ tumors were relatively immune-cold, while immune signatures became highly prominent in advanced malignancies. This involved changes in relative abundance of fibroblasts and immune cells, associated cell states and cell-cell interactions, which are consistent and synergistic with other reports^18,35,51^. Such changes also lead to rewired COL1A2 activities and extracellular matrix remodeling, especially in the advanced malignancies. Some other hallmarks^1^ such as altered cellular energetics arose early; we suspect that the selection for altered energetics became important to sustain clonal growth in a resource-limiting environment for tumors. Tumors can be viewed as dynamic complex systems, and from that viewpoint we took a note of the variations in the extent of changes in the cancer hallmark related biological processes within and between samples from different disease stages. We suspect that it showcases the functional heterogeneity of microenvironmental cues, and dysfunctional coordination thereof in increasingly malignant disease stages^62^. While Hu et al.^20^ examined the landscape of genetic changes in tissues from benign and malignant samples from lung, our findings provide a complementary and synergistic view of the multimodal changes in the tissue microenvironment states in the same samples.

Our effort to approach cancer as a complex, evolvable ecosystem was humbling, and we acknowledge several limitations to provide a balanced perspective. Single cell transcriptomics lacked the spatial and temporal contexts of disease progression. This is especially relevant while assessing intercellular interactions. Since many aspects of non-genetic changes in epithelial, immune, and stromal cell populations could be reversible and adaptive, analyses based on a limited cohort of tissue samples at a single time-point would need cautious interpretation. Tumor-immune interactions may themselves evolve in terms of modalities and complexities, which may not be fully captured here, given limited data and our focus on the global changes in the microenvironment. In addition, data on aging, lifestyle, environmental exposure, and treatment can have implications for immunity, health, and diseases, which in turn can influence the trajectory of tumorigenesis for the clonally amplified cell populations in individuals, as noted in the TracerX study^63^. Nonetheless, concordance between clonal makeups, transcriptional states and associated cancer hallmark-related biological processes, and histopathological classification suggest reliable and robust inference about functionally relevant changes captured here. However, OncoTerrain itself is not cancer-specific and the underlying architecture can be a resource for modeling the progressive chaos of the tissue microenvironment during tumorigenesis in other cancer types.

Our findings have broader significance. OncoTerrain was able to infer the composite pathway-level changes in the microenvironment at single cell resolution in increasingly malignant disease stages. It was especially relevant while analyzing the early-stage lesions, and we found that in situ carcinomas showed variation in the composite microenvironmental states that corroborated their histopathology, which was not apparent at the genome-level alone. The in-situ sample LT10 was a striking example, which was pathologically classified as an in-situ sample, but the microenvironmental signatures suggested potential for invasive growth that was corroborated in the detailed pathological notes. Abnormal pulmonary nodules are common in older adults, but a vast majority of suspicious pulmonary nodules do not progress to malignant disease^64^, such that concurrently improving early detection and minimizing over-diagnosis remain clinical priorities. While most malignant clonal growths are diagnosed as cancer, detection of benign lesions are often circumstantial findings, and the endpoints of their evolutionary trajectories remain unknowable. Our microenvironment-guided analysis suggests that often, such lesions may have a disrupted normal microenvironment that can facilitate tumor growth and associated hallmarks^1^. While genomic characterization of premalignant clones in the lung has so far offered some mechanistic insights, our analysis shows that many aspects of non-genetic changes accompanying, and often predating, the stages of cancer progression play important roles and need to be taken into consideration while classifying such cases.

## Methods

### Cohort summary

We obtained fresh frozen samples from benign pulmonary lesions (n=3), carcinoma in situ (n=5), and locally invasive or metastatic lung carcinoma (n=4), obtained from deidentified donors under an IRB approved protocol. Tumor adjacent pathologically normal tissues were procured from the cancer patients, while the benign pulmonary lesion samples were collected from preventive surgeries. Stains for infectious agents were negative.

### Single nuclei sequencing and data preprocessing

We performed single nuclei RNA sequencing of the snap-frozen tissue specimens using the 10X Chromium platform (Novogene), and then Cell Ranger was used to generate the initial expression matrix. Standard single-cell transcriptomic analysis was performed using Seurat^65^. After initial preprocessing and QC, we obtained 460-34736 unique barcodes per sample, and there were 267-1368 (25-75 percentile) transcripts expressed per barcode (**Figure 1B**). We primarily removed cells that had less than two hundred genes expressed and any genes that appeared in less than three cells. We further cleaned the data by removing any cells with above ten percent mitochondrial count and subsetting genes such that the counts per million (CPM) ranged from one hundred to six thousand. Log normalization was performed on the dataset and scaled with a factor of 10,000. Using variance stabilizing transformation, two thousand highly variable genes were obtained for downstream clustering analysis. We scaled log-normalized expression of the highly variable genes such that the mean and variance are 0 and 1, respectively.

### Single cell RNA sequencing cluster analysis and cell type annotation

We reduced the dimensionality using principal component analysis^66^ (PCA) and selected twenty dimensions by using an elbow plot to determine the significant clusters. We then constructed a k-nearest neighbors’ graph^67^ based on the Euclidean distance in PCA space and computed Jaccard similarity^68^. We utilized Leiden clustering^69^ to group cells together and choose a resolution of two as we had a large dataset; this resolution found the rare cell types and performed well. These clusters were tested using the silhouette coefficient^70^ to quantify the quality of the clusters. We further reduced dimensionality by running and visualizing uniform manifold approximation and projection (UMAP). UMAP^71^ was run with k = 10, 50, 100, and a minimum distance of 0.1, 0.2, 0.5. We found the optimal values of k and minimum distance to be 50 and 0.5 for this dataset. We identified marker genes by computing differential expression for each cluster. Then, those marker genes were subsetted to the top thirty based on -log(p-value) for further analysis. Those marker genes were cross-referenced with the Cell Marker Human database and the most frequent cell type for each cluster was chosen to be the annotation of the cluster. We utilized the molecular signature database^72^ (MsigDB) to find relevant genes associated with tumors and were used to show gene expression differences in different cell type populations.

### Celltype-specific transcriptional trajectory in pseudotime

We analyzed epithelial cells, myofibroblasts, fibroblasts, macrophages, and T cells, and identified further subclusters with cell-subtype-specific markers when applicable. The default monocle3 algorithm across all cell types was context-agnostic and linked unrelated cell types without any significant biological connection, motivating us to implement cell-type-specific trajectory inference based on a principal graph approach. For each non-epithelial cell type i.e. fibroblasts, macrophages, and T cells, we fit a principal graph to determine cell trajectory^36^. The trajectory was computed using the projection on the two principal components connecting the clusters using the minimal mean squared distance to the dataset. We further projected expression of selected marker genes on the cell-type specific barcodes to showcase the gradient in their gene expression along the respective trajectories.

### Pathway enrichment

We conducted the GSEA analysis^31^ by finding the differentially expressed genes in the cell-type specific clusters, and ranking the genes based on the log-norm fold change in the decreasing order. We obtained the cancer hallmark and immunologic gene sets from MsigDB and utilized Seurat’s built-in function AddModuleScore to calculate cell barcode-specific pathway scores. We further computed cell cluster-wise mean pathway scores at the sample-level by subsetting the cell barcode sets as appropriate and calculating the average score per barcode group.

### Ligand Receptor interactions and cell-cell communication

A disease stage-specific ligand-receptor analysis was conducted using CellChat^73^. with a study-objective specific modulation. We first subset the cellular barcodes according to disease stages and major cell types and grouped gene expression data accordingly. We then imported the human CellChat database and narrowed it down to relevant cell type subsets, which we assigned to each CellChat object. After ensuring that the cell identities were properly leveled, we computed the communication probabilities between cell types using the tri-mean method, filtered communications to include only those observed in at least ten cells, and computed pathway-specific communication probabilities. Finally, we aggregated the communication network, consolidating the interaction data to facilitate further analysis and interpretation of cellular communication within the cell population in the samples from a given disease stage. We visualized the cell-type to cell-type interaction patterns for various disease stages using a series of circos plots, detailing the interactions between different cell types for each signaling pathway. We then clustered the signaling pathways based on the expression of the genes therein; some pathways shared a subset of the genes or had similar biological contexts. The 2D projection of the pathways allowed us to visualize their potential synergy and coordination of the signaling processes in the tumor microenvironment from different stages of carcinogenesis in the lung.

### TabNet, our adaptations, and applications

We chose to use the TabNet classifier^74^ to capture synergistic multimodal signatures in different disease stages and to identify changes therein between the disease stages. We used it due to its sophisticated and efficient architecture tailored for handling tabular data. TabNet integrates a combination of decision trees and attention mechanisms, allowing it to selectively focus on relevant features and interactions while discarding less informative ones. The core of TabNet’s architecture is its attention-based decision steps, which dynamically highlight the most important features at each stage of the decision-making process. This feature selection is performed using sparse attention mechanisms, which not only improves the model’s performance by reducing noise but also enhances interpretability by emphasizing the most significant features. The model processes data through a series of decision trees, where each tree is built based on the attention scores from the previous step^75^. This iterative approach allows TabNet to capture complex feature interactions and hierarchical patterns that are crucial for making accurate predictions. We subsampled data from our cohort, when necessary, so that all cancer stages had similar amount of data points to prevent overfitting. We Min-Max scaled all the continuous features so that they were between -1 and 1 to further bolster performance of the model, and then performed a train-test split to set up our model. We performed a hyperparameter optimization for our TabNet using a randomized search approach. We defined a range of hyperparameters to explore and the search was conducted using StratifiedKFold cross-validation with 3 splits, ensuring that each fold maintains the proportion of classes in the training and validation sets. We tested all the classical ML metrics to understand the performance of our model. We utilized LT10 and LT2 for all testing purposes. The same steps used in our training data was performed on our testing cohorts and then outputted a UMAP of the histopathologist annotation vs the tuned-TabNet annotation of the cells to conduct in depth testing.

## Acknowledgement

The authors acknowledge scholarly input from other members of Rutgers Cancer Institute.

## Author contribution

V.V. designed experiments, performed analyses, interpreted the results, and wrote the manuscript with input from all authors. X.H. performed genomic analyses and interpreted the results. A.B. processed biospecimen and performed genomic analysis. A.S. provided guidance regarding genomic analysis. J.M. interpreted the results. G.R. provided pathological annotation and interpreted the results. S.D. conceived the idea, designed experiments, interpreted the results, and wrote the manuscript with input from all authors.

## Figures Information

**Figure 1: Overview of the study cohort and tissue composition**. A) Profiling transcriptomic landscapes of normal bronchia, benign lesions, in-situ carcinomas, invasive carcinomas, and metastatic carcinomas B) Location of sample and diagnosis from histopathologist C) Outline of study: Profiling the tissues, Cell type composition characterization with the onset of tumorigenesis, analyze cell-cell interactions, and utilize ML models to identify microenvironmental changes (D) Coloring the UMAP by sample ID and cancer stage (E) Coloring the UMAP by Cell Type Markers (F) Coloring the UMAP by Cell Type (G) Cell Type composition in the samples grouped by disease stages – i.e. benign lesions, in situ tumors, and invasion carcinomas.

**Figure 2: Functional heterogeneity of the epithelial cell population**. A) A schematic representation showing the clusters of epithelial cell populations are examined for functional characteristics and cancer hallmarks. B) Annotating the epithelial cell clusters on the UMAP based on epithelial cell subtype markers, activities of pathways related to known cancer driver genes and cancer hallmarks. (C) Boxplots showing changes in the activities of selected pathways related to cancer gene and hallmarks across different disease stages. Pathways associated with KRAS and EGFR activities, and apoptosis and epithelial mesenchymal transition are shown as examples. D) Relationship between subclones with distinct copy number differences and transcriptional heterogeneity in advanced carcinomas in lung. Sample LT14 and LT10 were shown as examples. E) Inferred transcriptional trajectories within the epithelial cell populations in LT14 and LT10 showing a gradient in transcriptional changes in pseudotime. F) Significant differences in selected cancer hallmark pathway activities between the sub-clonal clusters in LT14 and LT10. All between-cluster differences are significant (Mann Whitney U test, FDR adjusted p-value < e-02). F) Significant differences in selected cancer hallmark pathway activities between *EGFR*^*high*^*KRAS*^*high*^, *KRAS*^*high*^, and *other* samples in the cohort. All between-cluster differences are significant (Mann Whitney U test, FDR adjusted p-value < e-02).

**Figure 3: Functional heterogeneity of the non-tumor cell populations in the tissue microenvironment in the samples**. A) UMAP projections of the barcodes grouped by cell types and disease stages. Non-epithelial cells are shown by different colors, while epithelial cells are in grey. Within respective non-epithelial cell types, the barcodes from different samples show considerable overlap and stratification by disease stages. B) A schematic representation showing functional annotation and heterogeneity of the populations of non-epithelial cell types in the samples across different disease stages. C) Inferred transcriptional trajectories within the T cells in pseudotime and D) transcriptional gradient of selected genes along that transcriptional trajectory. E) Inferred transcriptional trajectories within the macrophages in pseudotime and F) transcriptional gradient of selected genes along that transcriptional trajectory. G) Inferred transcriptional trajectories within the fibroblasts in pseudotime and H) transcriptional gradient of selected genes along that transcriptional trajectory. I) Box plots showing differences in cell-type-specific COL1A2, STAT1, and CD163 expression in the samples that are EGFR^high^KRAS^high^, KRAS^high^, and *Other*. COL1A2, STAT1, and CD163 expression was shown in fibroblasts, T cells, and macrophages, respectively. J) Significant differences in the IL6-STAT pathway scores in the T cell populations between benign, in situ, and invasive carcinomas. Mann Whitney U test, FDR adjusted p-values < e-02. K) Significant differences in % fibroblast cells with detectable FAP expression between benign, in situ, and invasive carcinomas.

**Figure 4: Functional heterogeneity in intercellular signaling between stroma, immune, and epithelium in the samples**. A) Construct CellChat object to discover ligand-receptor interactions between tumor, immune, stroma and observe changes across disease stages. B) A dot plot showcasing the intensity of ligand-receptor interaction alterations through tumorigenesis, with x-axis representing cell type to cell type interaction and y-axis representing individual ligand-receptor pairs. C) A scatterplot visualizing signaling pathway interactions across disease stages, with each cluster representing closely interacting and/or related signaling pathways. (D) A 2D Heatmap showing correlation and expression of FN1-CD44 in benign, in-situ, and invasive samples. (E) A Sankey diagram showcasing incoming signaling pathways and outgoing signaling pathways for each pattern, where pattern was derived from NMF of the cellchat object. we have mapped each of those patterns to a corresponding cell type. (F) Circos plots showcasing Collagen, Laminin, and PTPRM signaling pathway augmentation through tumorigenesis, where we noted an increase in chaos consistent in many groups of intercellular interactions.

**Figure 5: Utilizing OncoTerrain to map changes in cancer hallmarks through disease stages and relate pathway-pathway activity in the samples**. A) Calculate barcode by pathway matrix to discover tumor state using microenvironment in different stages of disease. B) A reprojection of the barcodes using hallmark pathway module scores onto the UMAP hyperspace and we observe a far more homogeneous trajectory, albeit still heterogeneous, of tumor evolution. C) Coloring the UMAP with Allograft rejection pathway expression and we notice a correlation between intensity of expression and stage of cancer. D) Outline a methodology consisting of: (1) Feature Selection, (2) Curating a training set, (3) Tuning the hyperparameters of OncoTerrain (4) Inferencing upon individual samples. E) A 2D heatmap showing correlation and expression of Interferon Alpha-Interferon Gamma signaling through tumorigenesis. F) Testing our HP-Tuned Tab Net on LT10, where we notice the microenvironment was more invasive, one cluster was more benign and that aligned with the clonal data, and one cluster was more invasive like. G) Circular bar graph showing hallmark pathway activity in relation to the cancer hallmarks through stages of disease.

